# A Comprehensive Analysis Comparing Isotropic ADC to BOLD-fMRI: Sensitivity to Resting State Networks and Grey to White Matter Functional Connectivity

**DOI:** 10.64898/2026.07.02.736082

**Authors:** Jasmine Nguyen-Duc, Arthur Spencer, Tommaso Pavan, Inès de Riedmatten, Saina Asadi, Jean-Baptiste Perot, Ileana O Jelescu

**Affiliations:** Department of Radiology, Lausanne University Hospital (CHUV), Lausanne, Switzerland; Faculty of Biology and Medicine, University of Lausanne (UNIL), Lausanne, Switzerland

## Abstract

While Blood Oxygenation Level-Dependent (BOLD) fMRI remains the gold standard for mapping functional brain networks with MRI, its vascular origins inherently conflate haemodynamic effects with neural activity, limiting its sensitivity in white matter (WM) or its interpretation in neurovascular diseases. Apparent Diffusion Coefficient (ADC) fMRI offers an alternative, diffusion-based contrast that is theoretically more sensitive to neuromorphological coupling and therefore more specific to neuronal activation, though investigated primarily during task-based conditions. This study aimed to comprehensively evaluate the efficacy of isotropic ADC-fMRI in detecting established resting-state networks (RSNs) and to extend this methodology to the investigation of grey-to-white matter (GM-WM) functional connectivity.

Our analyses revealed a gradient of ADC detectability shaped by the degree of static functional cohesion and structural tethering of each network. The visual and somatomotor networks, being both highly segregated and strongly anchored to underlying structural pathways, yielded the most robust detection. The default mode network (DMN) and dorsal attention network (DAN) reached group-level significance but with lower effect sizes, and their detection proved fragile across analytical approaches. The frontoparietal network (FPN) and salience network (SAN), whose functional identity is defined by dynamic cross-network reconfiguration, did not reach significance. This gradient partially mirrors the established hierarchy of network segregation observed in BOLD, while further suggesting that ADC sensitivity depends on the structural grounding of each network.

Furthermore, ADC demonstrated superior sensitivity to GM-WM functional coupling compared to BOLD. GM-WM functional connectivity profiles derived from ADC were significantly more aligned with underlying structural WM architecture across subjects. Taken together, these findings position isotropic ADC-fMRI as a viable complementary modality to BOLD, offering a more direct window into the neural and structural foundations of brain connectivity.

## 1 Introduction

In brain research, it has become common practice to investigate functional magnetic resonance imaging (fMRI) timecourses in the absence of explicit tasks or stimuli. Even at rest, the human brain sustains baseline activity, driven by the continuous internal cognitive processing [1]. A perceptual task, such as reading this article, would interrupt such processes but would still involve many of the same brain regions that were active at rest [1].

Whether in task-active or resting-state (RS) conditions, Blood Oxygenation Level Dependent (BOLD)-fMRI can detect neural activity in spatially distinct regions via neurovascular coupling, enabling the identification of regions that exhibit synchronised temporal fluctuations [2–5]. Such patterns suggest these regions are functionally connected through spontaneous neural processes and constitute the fundamental architecture of resting-state networks (RSNs) [6–15]. Functional connectivity (FC) is commonly investigated to reveal these networks and is quantified by computing temporal correlations between BOLD signals from grey matter (GM) regions of interest (ROIs).

Substantial research effort has been directed towards examining FC between GM ROIs. However, BOLD signals directly originating in WM are rarely investigated. Often, they are considered physiological noise, thus treated as a covariate and regressed out from the BOLD signal [16]. Nonetheless, there has recently been growing interest in using fMRI to detect brain activity in WM. Converging evidence indicates that meaningful information related to interregional communication could be inferred from WM BOLD signals, and should therefore not be overlooked [17–27]. Integrating WM FC analysis could help uncover specific tracts that mediate information transfer between GM regions in different conditions [17]. Such an analysis could deepen our understanding of brain connectivity and may also help differentiate healthy individuals from those with neurodegenerative and neuropsychiatric conditions at early stages of disease [19, 23, 24].

However, although BOLD signals in WM could potentially contain functional information, their detection and interpretation is complicated by a sparse vascular architecture [28] and a haemodynamic response function (HRF) that deviates significantly from GM [29]. Traditionally, WM metabolic costs have been attributed to glial maintenance [30]. Some studies [25] suggest that glial functions, including ion sequestration and nutrient provision, generate a measurable (albeit distinct) HRF. However, recent evidence from the optic nerve suggests that axonal activity may instead be fuelled by internal oligodendrocyte-to-axon metabolic loops that bypass neurovascular coupling entirely [31]. If WM lacks a locally-driven, activity-dependent HRF, the variable signals often detected may reflect non-signalling glial processes or vascular spillover rather than neural activity. Such physiological ambiguity continues to make the inclusion of WM in FC analyses a point of considerable debate.

To circumvent the limitations and ambiguities related to BOLD-fMRI, studies have investigated alternative fMRI contrasts based on diffusion MRI. Because diffusion MRI reflects the interaction of water molecules with physical obstacles, cellular swelling and microstructural deformations associated with neural activity [32] are believed to drive the overall decrease in measured water diffusion. This neuromorphological coupling was investigated using diffusion weighted (DW) timeseries at specific b-values [33–37], and the apparent diffusion coefficient (ADC)-fMRI [36–44]. The latter has been shown to remain independent of vascular contributions [45], whereas conventional DW signals are notably confounded by these effects [46].

In RS, ADC-fMRI was able to identify more significant GM–WM and WM–WM connections than BOLD in graph analyses [41], suggesting that ADC timecourses may be more sensitive to WM activity than BOLD. This previous work was conducted using linear diffusion-encoding ADC-fMRI, which yields sensitivity to cell morphological changes in a single specific direction, as similarly done by [42] in task. This can introduce inhomogeneous sensitivity to functional ADC changes throughout the WM, as the sensitivity is maximal perpendicularly to the WM fibres [39, 43]. Recent developments in ADC-fMRI protocols now incorporate isotropic diffusion encoding [47] which sensitises the signal to diffusion in all directions of space in a single shot. Isotropic ADC-fMRI thus successfully captures morphological fluctuations regardless of the geometric orientation of the WM tracts [43, 44]. Additionally, previous RS ADC-fMRI work [41] relied on a limited spatial coverage during the acquisition, limiting the analysis to a brain slab. For this reason, large-scale RSNs could not be assessed.

With this work, we tackled these limitations, comparing isotropic ADC-with BOLD-fMRI using whole-brain (excluding cerebellum) RS data. The first aim was to identify how the RSNs revealed by ADC-fMRI compare to those typically reported in BOLD-fMRI. This was done using the following methods: Independent Component Analysis (ICA) [5, 12, 48–50], Seed-based Analysis [5, 8, 11, 13, 51–53] and ROI-ROI FC with graph analysis [54–56, 56–59]. In the latter, RSN detection was assessed by measuring the segregation of each RSN. Our second aim was to investigate each modality’s sensitivity to GM-WM functional coupling. Although the relationship between FC and structural connectivity (SC) is extensively studied [55, 60], discrepancies remain heavily debated, as standard BOLD-fMRI frequently detects functional correlations between brain regions that lack direct anatomical connections. We hypothesised that ADC-fMRI, by bypassing vascular confounds, would yield GM-WM FC more closely aligned with underlying SC, which would suggest that these mismatches largely stem from BOLD-related vascular blurring rather than a true divergence between neural activity and structural pathways.

## 2 Methods

### 2.1 Acquisition

This study was approved by the ethics committee of the canton of Vaud, Switzerland (CER-VD). All participants provided written informed consent. Fifteen healthy volunteers participated and completed two resting-state runs in a single session, one using ADC-fMRI and one using BOLD-fMRI. Each resting-state run lasted 14 min 40 s, during which participants were instructed to fixate on a cross. MRI data were acquired as described in [43, 45] on a 3T Siemens Magnetom Prisma system equipped with 80 mT/m gradients, a slew rate of 200 T/m/s, and a 64-channel head coil.

- **High-resolution whole-brain** *T*_1_ **anatomical images** were obtained for anatomical reference and tissue segmentation using a 3D Magnetization Prepared 2 Rapid Acquisition Gradient Echoes (MP2RAGE) sequence [61]. Acquisition parameters were as follows: 1 mm^3^ isotropic voxels; repetition time (TR) of 5000 ms; echo time (TE) of 2.98 ms; inversion times (TI) of 700 and 2500 ms; flip angles of 4° and 5°; and an in-plane acceleration factor of 3 using Generalized Autocalibrating Partially Parallel Acquisitions (GRAPPA) [62].
- **Diffusion fMRI data** were acquired using a diffusion-weighted spin-echo EPI sequence with spherical diffusion encoding, employing a waveform compensated for cross-terms with background field gradients [47]. The diffusion waveform was generated using the NOW toolbox (https://github.com/jsjol/NOW). Acquisition parameters included: TE/TR = 105/1100 ms; in-plane resolution: 2.8 mm^2^; matrix size: 82 x 82; field of view: 232 × 232 mm^2^; 24 2.8-mm thick slices with a 40% slice gap; GRAPPA acceleration 2; partial Fourier factor of 6/8; multiband acceleration factor 3. Each diffusion fMRI run included two initial b = 0 volumes, followed by alternating volumes with b-values of 200 and 1000 s/mm^2^. This resulted in two timeseries: dfmri200 and dfmri1000. These timeseries were necessary to compute the ADC-fMRI timeseries. Two additional b = 0 s/mm^2^ volumes with reversed EPI phase encoding direction were acquired to enable correction of field inhomogeneity distortions.
- **Multi-echo BOLD-fMRI data** [63] were acquired with a gradient echo EPI sequence (TE 12.60, 30.18, 47.76, 65.34 ms) with a flip angle of 62°. All resolution and acceleration parameters were matched to the ADC-fMRI acquisitions. Two additional volumes were acquired with reversed EPI phase encoding direction for correction of field inhomogeneity distortions.

### 2.2 Preprocessing

Data were preprocessed according to [43, 45], as follows:

- **For** *T*_1_**-weighted data** Denoising was done using spatially adaptive non-local means filtering with ANTs [64], then skull-stripped with Synthstrip [65].
- **For diffusion fMRI data**, magnitude image denoising was applied to dfmri200 and dfmri1000 timeseries separately using NORDIC [66], with a 7 x 7 x 7 kernel and step size 1 for both g-factor estimation and PCA denoising. Gibbs unringing was applied to all volumes using MRtrix3 [67, 68]. The susceptibility off-resonance field was estimated from b = 0 volumes, and subsequently used to correct susceptibility-induced distortions in all volumes, using FSL Topup [69, 70]. Both timeseries were corrected for motion using ANTs [71], then registered to the initial b = 0 volume. Absolute displacement from this first volume and framewise displacement between volumes [72] were calculated from motion correction parameters for dfmri200 and dfmri1000 and runs with a maximum absolute displacement greater than one voxel or a mean framewise displacement greater than 0.2 mm were rejected. In total, dfMRI runs from two subjects were rejected due to excessive movement, resulting in n = 13. A brain tissue mask was created from the Topup-corrected b = 0 volumes using Synthstrip [65] and used to remove non-brain voxels from the full timeseries.
- **For BOLD-fMRI data**, motion correction parameters were calculated on the timeseries of the first echo, then these transformations were used to correct the timeseries for all echoes, using ANTs. As for diffusion fMRI, runs with a maximum absolute displacement greater than one voxel or a mean framewise displacement greater than 0.2 mm were rejected. Two RS runs were rejected due to excessive movement, resulting in n = 13. One of the rejected runs was shared with dfMRI. An optimally combined signal was calculated from a weighted average of echoes using Tedana, including denoising with TEDPCA and TEDICA [73]. FSL Topup was applied to the first echo to calculate the susceptibility off-resonance field, then this was used to correct distortions in the optimally combined data. A brain mask was created using Synthstrip to remove non-brain voxels.
- **ICA cleaning** was used to remove physiological noise separately from each timeseries (dfmri200, dfmri1000, and BOLD) by regressing out selected independent components with FSL’s MELODIC ICA. For each subject and timeseries, 100 components were extracted and manually classified as either signal (if associated with neural activity) or noise (if associated with ventricular signal, motion-related edge effects, slice cross-talk, or fat artefacts). Each class was identified based on characteristic features observed in the spatial maps of the components. For instance, components were labelled as CSF/ventricular when high z-scores were predominantly located within the CSF, and as slice cross-talk when elevated z-scores were confined to slices acquired simultaneously. Motion-related components were characterised by high z-scores along the brain boundaries, whereas fat artefacts exhibited ring-like spatial patterns and a prominent spectral peak around 0.13 Hz. Slice cross-talk and fat artefacts were only present in dfmri200 and dfmri1000 timeseries. Supplementary Figure S1 shows how many components were classified in each category. In total, approximately 58%, 36% and 55% of components were regressed out for dfmri200, dfmri1000 and BOLD timeseries respectively. Supplementary Figure S2 shows see spatial maps for each category of regressed component in an example subject.

### 2.3 Creating ADC timeseries

The ADC-fMRI timeseries was calculated from preprocessed dfMRI data using the following equation:

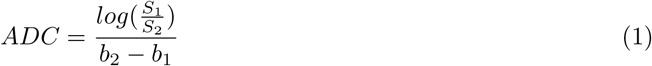

where *S*_1_ corresponds to dfmri200, *S*_2_ to dfmri1000, and *b*_1_ and *b*_2_ denote the corresponding b-values of 200 and 1000 s/mm^2^ respectively. Computing the ADC timeseries in this manner minimises *T*_2_ weighting and direct intravascular signal contributions, thereby reducing BOLD contamination.

### 2.4 GM and WM ROI Parcellation

#### 2.4.1 GM ROIs

To investigate the presence of RSNs within both BOLD and ADC modalities, we utilised the Schaefer functional atlas consisting of 200 GM ROIs, which was registered to subject space using ANTs. This atlas is segmented into ROIs that have each been associated with an RSN. For quality control, the left and right ‘Vis5’ areas were excluded as their voxel counts fell below the minimum threshold of 10 (in subject space), resulting in a final set of 198 ROIs. These regions were categorised into seven canonical RSNs based on the Yeo et al. (2011) parcellation [14]: the Default Mode (DMN), Dorsal Attention (DAN), Salience (SAN), Frontoparietal (FPN), Somatomotor (SMN), Visual (VIS), and Limbic (LIM) networks. However, only cortical regions were included in this atlas, despite the LIM network being mostly composed of subcortical regions. For this reason, the LIM network was overlooked for this study.

In the analysis on the GM-WM connectivity, GM ROIs were defined using the Desikan–Killiany (DK) anatomical atlas [74] provided with FreeSurfer. From the resulting 84 regions, we excluded the cerebellum, accumbens area, and entorhinal cortex because these ROIs were either outside the field of view or did not meet our minimum 10 voxel inclusion criterion in subject space. Following these exclusions, 78 GM ROIs were retained for analysis. The DK atlas was selected over the Schaefer functional parcellation to ensure spatial correspondence with the *probconnatlas* tool, which was used to create WM ROIs in subsequent processing stages.

#### 2.4.2 WM ROIs

The *probconnatlas* tool [75] provided spatial probability maps of WM pathways connecting each pair of cortical ROIs from the DK atlas, based on tractography from 66 HCP subjects. This yielded over 4,000 pairwise tract probability maps which were binarised. In order to reduce the number of maps, only those that were composed of over 100 non-zero voxels (in MNI standard space) were considered. They were then arranged in a hierarchical tree, where the distance metric between two maps was defined as 1 - the Dice score, and therefore varies inversely with the degree of spatial overlap. Consequently, maps that are closer to one another in the tree exhibit greater spatial overlap. This hierarchical clustering was performed separately on three sets of WM tracts: (i) tracts connecting pairs of cortical GM ROIs within the right hemisphere, (ii) tracts connecting subcortical and cortical GM ROIs within the right hemisphere, and (iii) tracts connecting cortical GM ROIs across hemispheres, from left to right. For cases (i) and (ii), the corresponding left–left WM tracts were generated by applying the same ROI–ROI clustering to the left hemisphere. This procedure yielded three tract classes: (i) 52 association fibres (26 per hemisphere), (ii) 24 projection fibres (12 per hemisphere), and (iii) 14 commissural fibres. In total, this resulted in 90 WM fibre tracts. Examples of these fibres can be visualised in Supplementary Figure S3. Following non-linear registration to subject space using ANTs [71], each WM tract was strictly constrained using a subject-specific WM mask derived from the individual’s *T*_1_-weighted structural image. It is important to keep in mind that these WM tracts were not derived using subject-specific tractography, but were instead transformed into subject space from MNI space.

### 2.5 Independent Component Analysis

Group-ICA was used to extract RSNs, with 100 components selected for BOLD. For ADC-fMRI, we allowed MELODIC FSL to automatically estimate the dimensionality, resulting in 250 components, since this is the first study to apply ICA to this contrast and no prior guidance on an appropriate model order exists. This allowed for an explained variance of approximately 55% for both modalities. We then compared the spatial maps obtained to identify how the RSNs revealed by ADC-fMRI compare to those typically reported in BOLD-fMRI.

### 2.6 Seed-based Analysis

Seed-based analyses were conducted using FSL FEAT with mixed-effects modelling (FLAME 1+2). Following the Schaefer atlas [76], representative seeds were chosen for six established RSNs: DMN (precuneus/posterior cingulate), FPN (lateral prefrontal cortex), SMN (precentral gyrus), VIS (lateral occipital cortex), DAN (superior parietal cortex), and SAN (anterior insula/operculum). We analysed left and right hemisphere seeds independently. Statistical maps were cluster-thresholded (*z >* 2.3, *p <* 0.05) to identify significant voxels. We assumed that ROI seeds are functionally connected to other ROIs within the same RSN. Results with and without Global Signal Regression (GSR) were investigated.

### 2.7 ROI-ROI FC Graph Analysis

Graph analysis has been applied in two separate analyses: the segregation of RSNs and the GM-WM connectivity. The former was done with Schaefer parcellation whereas the latter was done with the DK atlas.

In both analyses, FC graphs using ROI-ROI connectivity were computed for each subject and contrast. A Pearson’s correlation matrix was measured in subject space and Fisher z-transformed. Only positive correlations were retained. One-sample t-tests corrected for multiple comparisons (FDR, *α* = 0.01) were used to keep correlations that agreed across subjects. To enable thresholding, each subject’s matrix of (positive) Fisher z-transformed correlations was then standardised (z-scored) across all edges, yielding a relative connection-strength z-score for each edge. This standardised z, rather than the Fisher z-transformed correlation itself, is what is referenced when matrices are binarised across thresholds. Note that it reflects each connection’s relative strength within the subject’s own connectivity distribution, and should not be confused with the Fisher z-transformation applied earlier.

#### 2.7.1 Resting State Network segregation using the Schaefer atlas

A fundamental defining characteristic of any RSN is its functional segregation: on average, nodes are expected to communicate more densely within their designated network than with external cortical regions. Although previous research quantified segregation using the Participation Coefficient, which evaluates connectivity globally across all brain modules [54, 77, 78], this metric does not isolate intra-network correspondence specifically. We therefore calculated the within-module connection fraction, which directly measures whether nodes in each RSN communicate preferentially within their own module. For a given node *i* in module *m*, this is computed as:

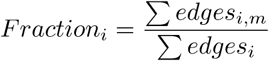

where Σ*edges_i,m_* denotes the number of edges between node *i* and all other nodes specifically within its module *m*, and the denominator Σ*edges_i_* represents the total degree of node *i* across the entire graph. This nodal metric was averaged across all nodes within each RSN and evaluated within FC matrices thresholded at *z >* 2.3, as illustrated in Figure 1A.

**Figure 1:**
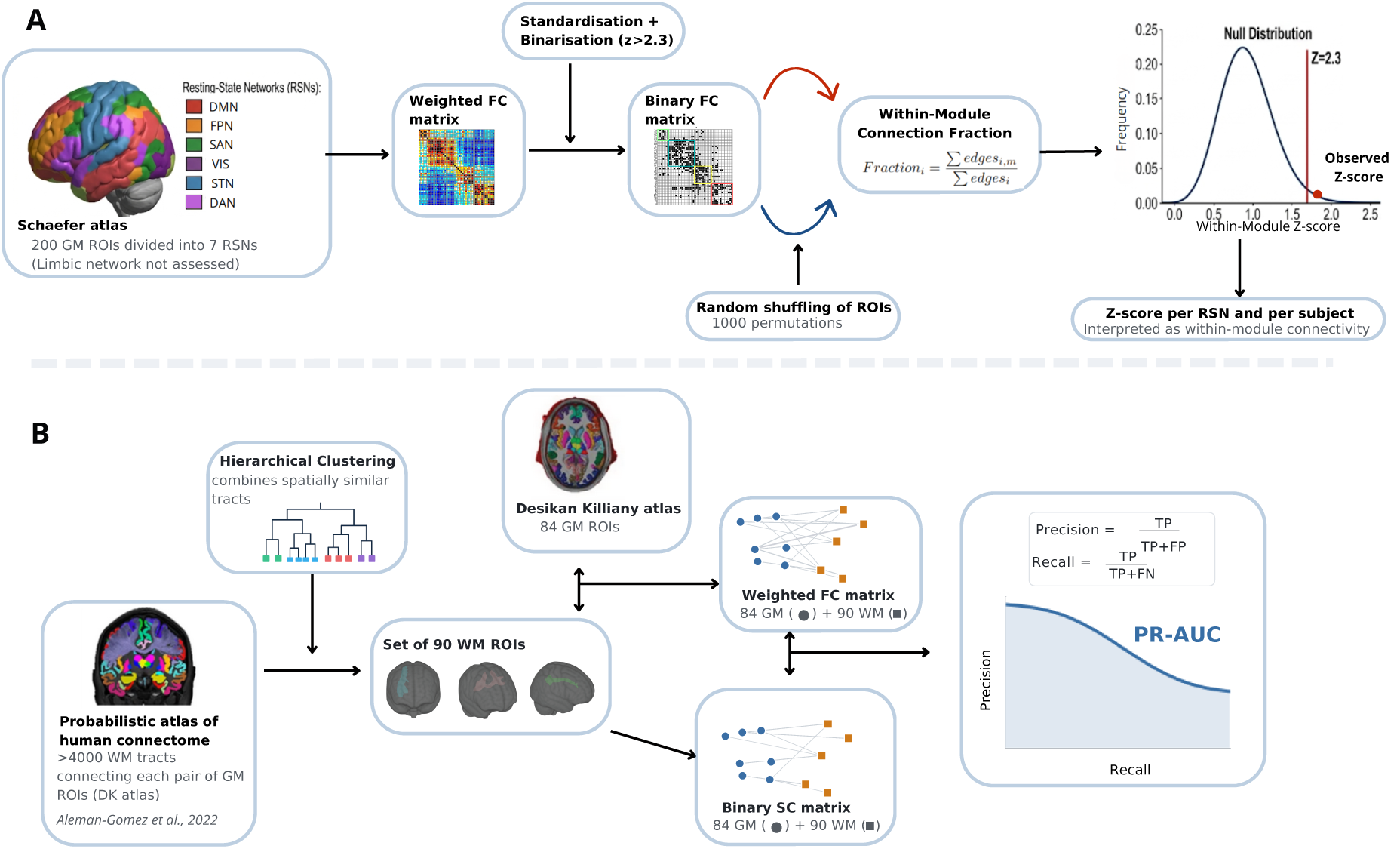
Pipeline for ROI-ROI graph analysis. (A) GM–GM functional connectivity (FC) is assessed using the Schaefer atlas, where each of the 200 ROIs is assigned to one of seven resting-state networks (RSNs) as defined by [14]. For each subject, a FC matrix is derived from Fisher-transformed Pearson correlations, then standardised and binarised using a threshold of *z >* 2.3. The within-module connection fraction is computed for each RSN (excluding the LIM network) and compared against its corresponding null distribution, yielding a z-score per RSN, subject, and modality. (B) The *probconnatlas* tool [75] is used to generate WM tracts connecting each pair of GM ROIs defined by the DK atlas. The number of WM tracts is reduced to 90 ROIs by merging spatially overlapping tracts via hierarchical clustering. A weighted FC matrix is then computed, restricted to GM–WM connections, and evaluated against the binary SC matrix to derive a PR-AUC score per subject and modality.

To assess whether the observed segregation exceeded chance, we compared each network’s empirical average against a null distribution derived from 1000 node label permutations per RSN, subject, and contrast. This permutation strategy preserved the fundamental topology of the graph (including exact edge weights and degree distributions) while entirely disrupting the assignment of anatomical regions to functional modules. Raw intra-module connection fractions are sensitive to variations in module size and network density, so each empirical value was standardised against its null distribution to yield a *z*-score, controlling for these topological biases and enabling direct comparison across subjects and modalities. A *z*-score above 2.3 was considered significant at the individual subject level, and group-level significance was then assessed using a one-sample t-test of the subject-wise *z*-scores against this threshold, with FDR correction (*α <* 0.01) across the six networks.

#### 2.7.2 GM-WM structure-function coupling

FC graphs were designed with both WM and GM as nodes, using the *probconnatlas* and the DK atlas respectively. Following subject-wise matrix binarisation using multiple thresholds (0 to 3.1), we compared the number of significant connections for each type (categorised as GM–GM, WM–GM, or WM–WM) between the two modalities.

Structure-function coupling was then investigated by comparing the weighted GM-WM FC with a binary SC matrix, as illustrated in Figure 1B. The SC matrix was constructed by identifying the GM ROIs connected to each tract, traced from the underlying WM pathways. For FC, all GM-GM and WM-WM connections were discarded before the standardisation and binarisation of matrices, thus keeping only the GM-WM connections. The agreement between FC and SC was quantified using the Precision-Recall Area Under the Curve (PR-AUC), which was computed per subject and per contrast.

More specifically, the correspondence between SC and FC was evaluated by iteratively binarising the FC matrices across a range of thresholds (*z* = −3.1 to 3.1). At each threshold, precision and recall metrics were calculated relative to the binary SC matrix, allowing for the construction of a precision-recall (PR) curve. The precision and recall metrics can be computed as follows:

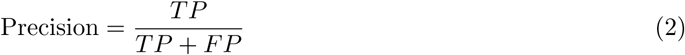

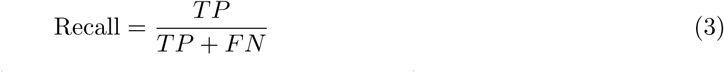

where TP represents true positives (positive instances correctly identified), FP denotes false positives (negative instances incorrectly labelled as positive), and FN stands for false negatives (positive instances incorrectly missed). The PR-AUC was computed for each individual matrix to provide a single-value estimate of SC–FC agreement per subject and per contrast.

## 3 Results

### 3.1 Independent Component Analysis

The independent components derived from BOLD data revealed spatial maps corresponding to RSNs, such as the DMN, SAN, and VIS networks. Figure 2 shows representative examples of these BOLD components. For each BOLD component, the independent ADC components that exhibited the greatest spatial overlap are shown. In these examples, the spatial correspondence between BOLD and ADC components is high.

**Figure 2:**
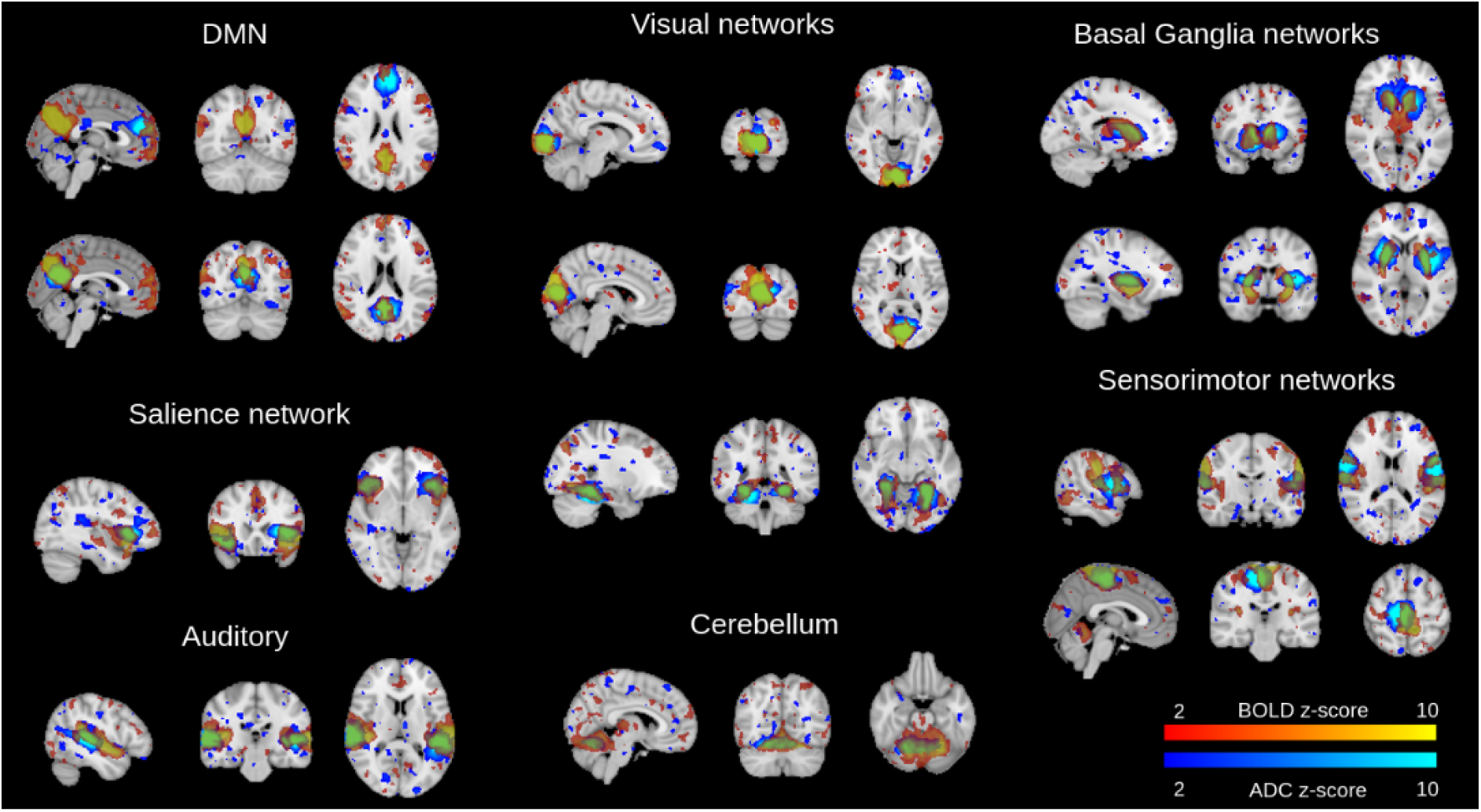
Group-level ICA was performed using MELODIC FSL. Representative ADC components (blue) are overlaid with their corresponding BOLD counterparts (orange). Because ADC RSNs were spatially fragmented across multiple independent components, several were manually combined prior to comparison with BOLD.

It is important to note, however, that RSNs comprising spatially disconnected ROIs could not be captured within a single ADC-fMRI component. Even for bilateral ROIs, one hemisphere typically appeared in one component and the contralateral hemisphere in another. Consequently, in Figure 2, a single BOLD component is often shown alongside several combined ADC components on the same brain.

### 3.2 Seed-based Analysis

Our analysis utilised a seed-based approach applied to six ROIs from each hemisphere (12 in total), with each region selected to represent a different RSN. The results are shown without GSR (Figure 3) and with GSR (Supplementary Figure S4).

**Figure 3:**
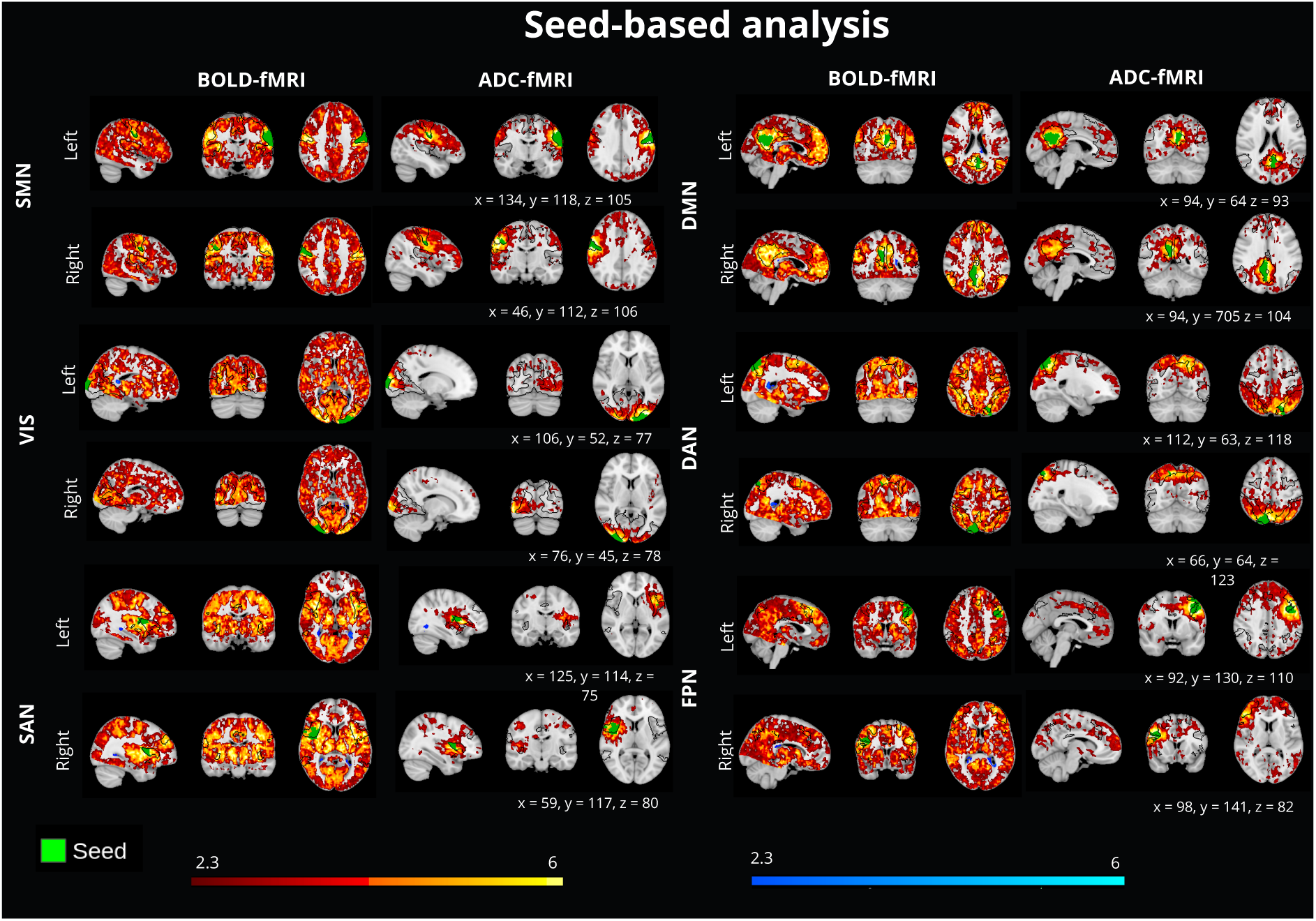
Group-level seed-based analysis performed on seeds including: the precuneus/posterior cingulate (DMN), lateral prefrontal cortex (FPN), precentral gyrus (SMN), lateral occipital cortex (VIS), superior parietal cortex (DAN), and anterior insula/operculum (SAN). Both left and right ROIs were taken as seeds in separate maps. Statistical maps were obtained using FSL FEAT with mixed-effects modelling (FLAME 1+2) and thresholded at an absolute *z >* 2.3 (cluster-corrected) to identify significant voxels.

In the BOLD analysis, without GSR, an extensive spatial coverage of significant voxels could be noticed for all seeds, with overall much higher statistical significance (z-scores) than ADC-fMRI. The permissive threshold of z=2.3, which was selected to ensure a direct comparison with the ADC results, allowed for voxels to be significant even if they were not within the RSN of interest. RSNs remained discernible across most maps, characterised by hyper-intensities that were consistent with the seed’s respective network.

In the ADC-fMRI analysis, the voxels with the highest z-scores were localised within the seed regions and their immediate spatial vicinity. Nonetheless, ADC sometimes captured significant clusters that were spatially disconnected and far away from the seed. Some of these clusters seemed to align with the contralateral ROI and extended to some parts of the associated RSNs. For example, seeding the precuneus within the DMN revealed functional coupling with the medial prefrontal cortex and the angular gyri. However, some significant clusters of similar intensity were also observed in unexpected regions such as the occipital cortex, indicating the potential presence of false positives.

Regarding the application of GSR, it markedly attenuated the spatial extent of significant positive BOLD connectivity compared to the uncorrected data. Furthermore, significant negative z-scores clusters appeared in distinct regions. For ADC, applying GSR did not substantially alter the spatial extent of the positive clusters, though it suppressed most positive regions situated distally from the seed. However, ADC maintained some bilateral connectivity across several RSNs, most notably in the SMN and VIS networks. While the regression procedure also induced the appearance of new negative connectivity clusters, these were generally small and spatially scattered. Furthermore, seeding specific ROIs in the ADC data induced sparse but significant negative correlations that spatially mirrored those observed in BOLD. For instance, when utilising the precuneus (DMN) as a seed, both modalities detected negative functional coupling within the precentral gyrus. Notably, while BOLD captured this anti-correlation bilaterally, the ADC signal was spatially restricted to the right precentral gyrus.

### 3.3 Graph Analysis

#### 3.3.1 GM Resting-State Networks

Group-averaged connectivity matrices (Figure 4A–B) were derived from subject-level binary matrices. A divergence was observed in interhemispheric connectivity, where BOLD-fMRI exhibited strong coupling that was largely absent in ADC.

**Figure 4:**
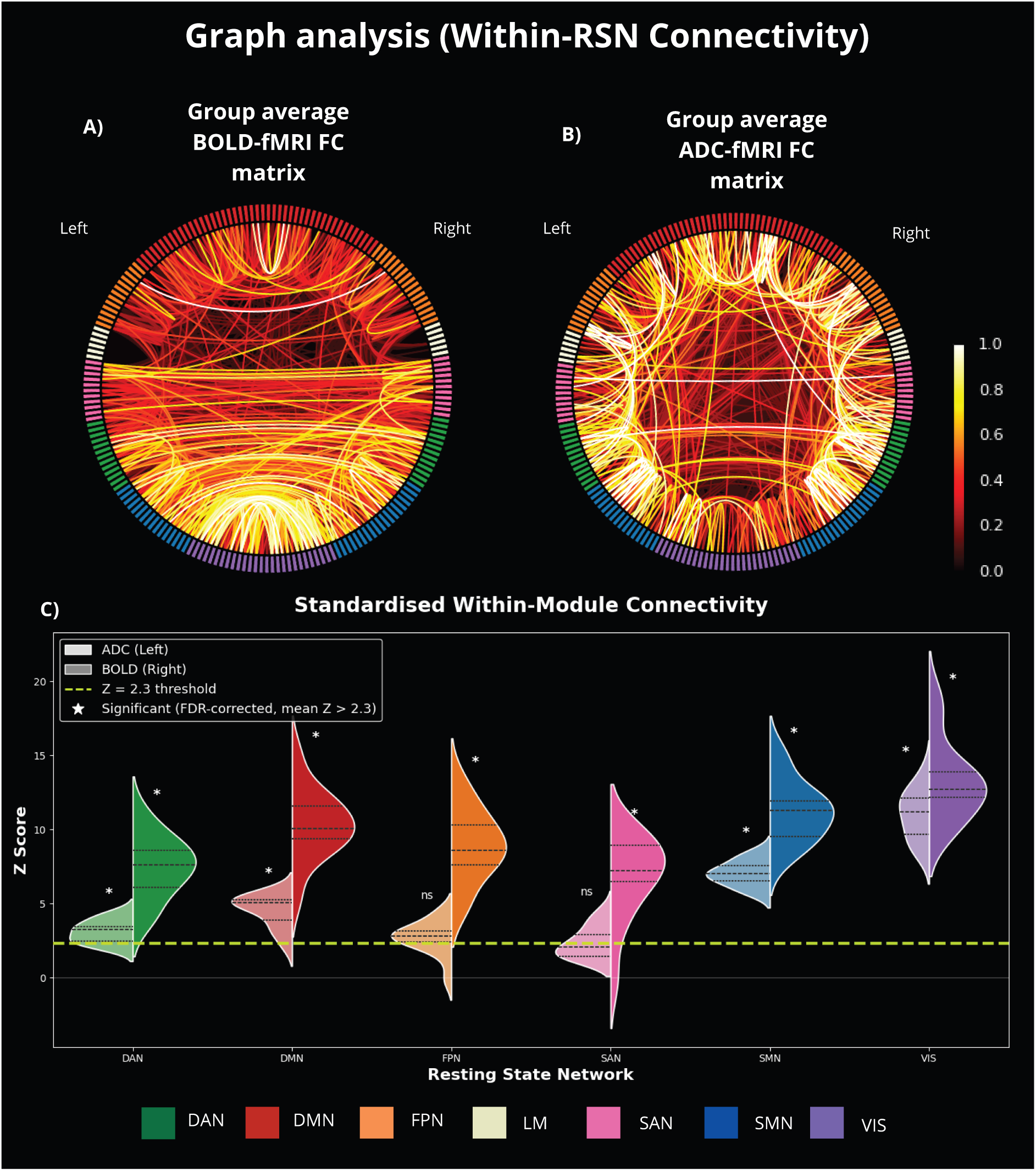
Functional Connectivity Group-level consensus matrices for BOLD- (A) and ADC-fMRI (B). All ROIs from each RSN are shown, including those within the Limbic network (LIM) though this RSN is not included in the subsequent analysis. Individual ROI-ROI matrices were computed using Fisher-transformed Pearson correlations. Edges inconsistent across subjects were discarded (one-sample t-test, FDR *α <* 0.01), and surviving connections were binarised at a strength threshold of *z >* 2.3. Intensity represents the percentage of subjects possessing a significant connection at each edge. (C) Statistical validation of RSN segregation. All RSNs are investigated except for the LIM network due to the absence of subcortical GM in the atlas. The within-module connectivity fraction was computed for each node and averaged per RSN. This average was then compared against a label-shuffling null distribution (1000 permutations) to generate a subject-specific Z-score, where higher values indicate stronger functional segregation. Group-level significance was assessed using a one-sample t-test of subject-wise Z-scores against z=2.3 (FDR-corrected, *α <* 0.01), with * denoting significant networks and ns denoting non-significant ones. The green dashed line marks the z=2.3 reference threshold. Inter-subject variability is shown in split violin plots, with dashed lines indicating the 25th percentile, median, and 75th percentile.

A within-module connectivity fraction was calculated for each RSN per subject and modality. *z*-scores were derived by comparing the observed data against a null distribution, where higher values indicate stronger functional segregation. As depicted in Figure 4C, within-module connectivity fractions were substantially larger for the BOLD timecourses compared to ADC across all evaluated networks. Within the BOLD data, the DMN, VIS, and SMN networks exhibited exceptionally high segregation, with an average *z*-score above 10. All remaining networks evaluated via BOLD yielded *z*-scores significantly greater than 2.3 (one-sample t-test, FDR-corrected, *α <* 0.01). While the ADC modality yielded consistently lower absolute segregation values, it preserved a similar hierarchical network order. The VIS, SMN, DMN, and DAN each displayed group-level segregation significantly exceeding *z* = 2.3(one-sample t-test, FDR-corrected, *α <* 0.01), with the VIS and SMN yielding the highest absolute values. The FPN and SAN did not reach group-level significance, with more than 25% of subjects recording *z*-scores below 2.3 for these networks.

#### 3.3.2 GM–WM Structural-Function Coupling

An analysis of the number of significant connections divided by tissue type revealed distinct functional profiles for each modality. As illustrated in Figure 5A, BOLD-fMRI consistently yielded a greater number of GM-GM connections compared to ADC-fMRI across all evaluated Z thresholds. Conversely, ADC-fMRI demonstrated a superior yield of WM-WM connections specifically at statistical thresholds exceeding *Z >* 1. Furthermore, across the entire evaluated spectrum of z-scores, the GM-WM connections remained persistently higher in the ADC modality than in BOLD.

**Figure 5:**
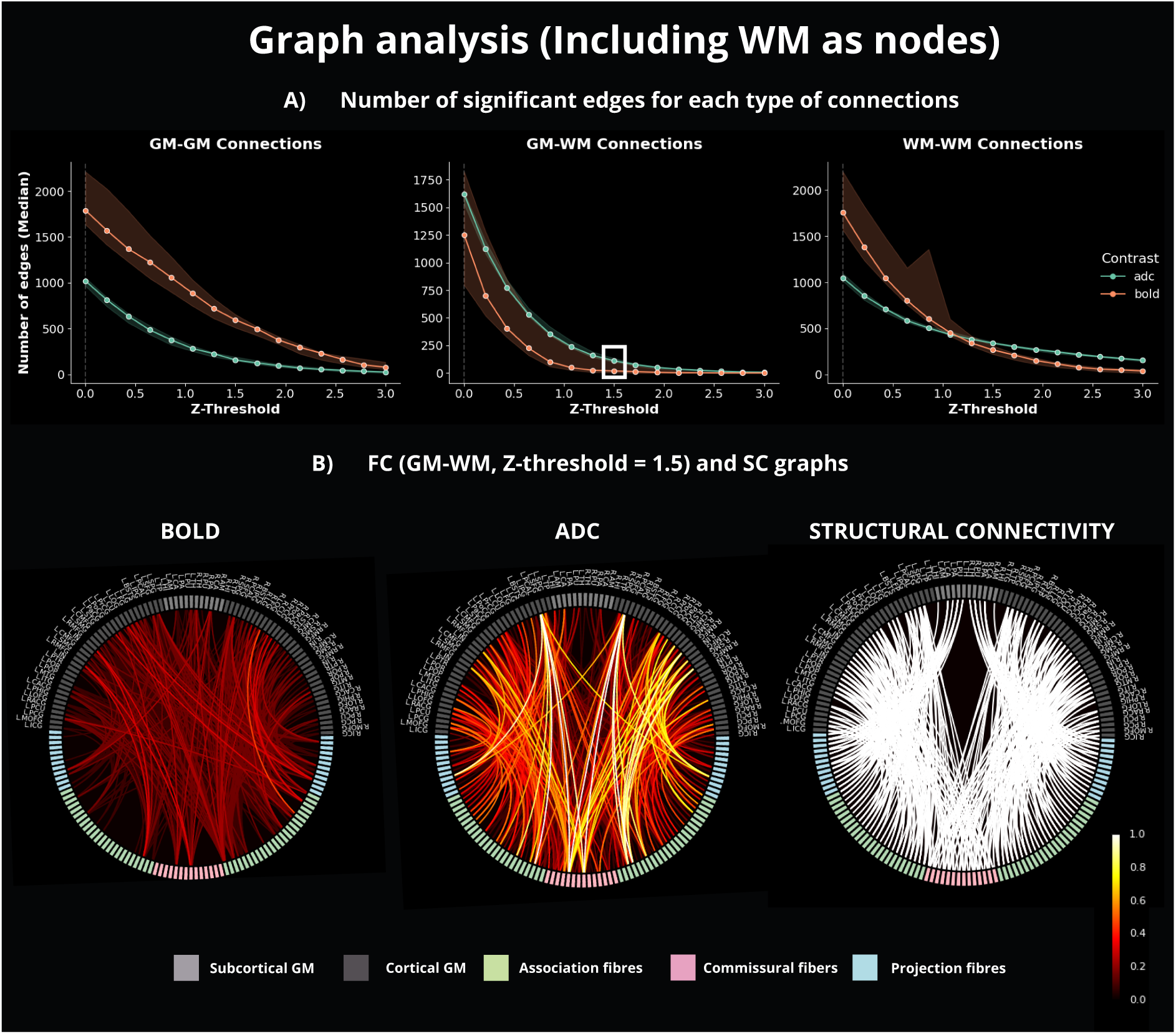
(A) Number of edges in FC matrix connecting: GM-GM, GM-WM and WM-WM. This is shown when varying the z-threshold. (B) Group-average FC graphs including only GM-WM connections, obtained from subject-specific binary matrices thresholded at *z >* 1.5. Results are shown for BOLD, ADC and structural connectivity. The latter was not subject-specific and was obtained using the *probconnatlas* tool. The scale indicates the proportion of subjects for which the edge was significant.

Figure 5B illustrates the group level correlation matrices isolating the strongest GM-WM connections, defined by a statistical threshold of *Z >* 1.5. These matrices were aggregated from subject level binarised data, where higher matrix intensities reflect a specific connection reaching statistical significance across a larger proportion of the cohort. Evaluating the BOLD-fMRI data revealed a dichotomous pattern as numerous WM regions demonstrated no significant connectivity to any GM targets. Conversely, this all or nothing topology was absent in the ADC-fMRI data, where every individual WM tract maintained a significant connection to at least one GM region. Furthermore, the ADC matrices exhibited elevated overall intensities, demonstrating that significant GM-WM coupling was detected in a larger fraction of subjects compared to the BOLD modality. To provide an anatomical reference, the binary SC matrix is additionally presented in Figure 5B, facilitating a direct visual comparison against the FC matrices.

The PR-AUC analysis in Figure 6 revealed that the ADC-fMRI graph aligned more closely with the binary SC graph than the BOLD-fMRI graph: ADC yielded significantly higher PR-AUC values than BOLD (median value at 0.270 vs 0.105, Wilcoxon signed-rank test, paired, two-sided; *p* = 2.4 *×* 10*^−^*^4^), indicating a better correspondence between SC and ADC-based FC. As previously described, SC in this context simply maps each WM tract to each of the GM ROIs it is structurally connected to.

**Figure 6:**
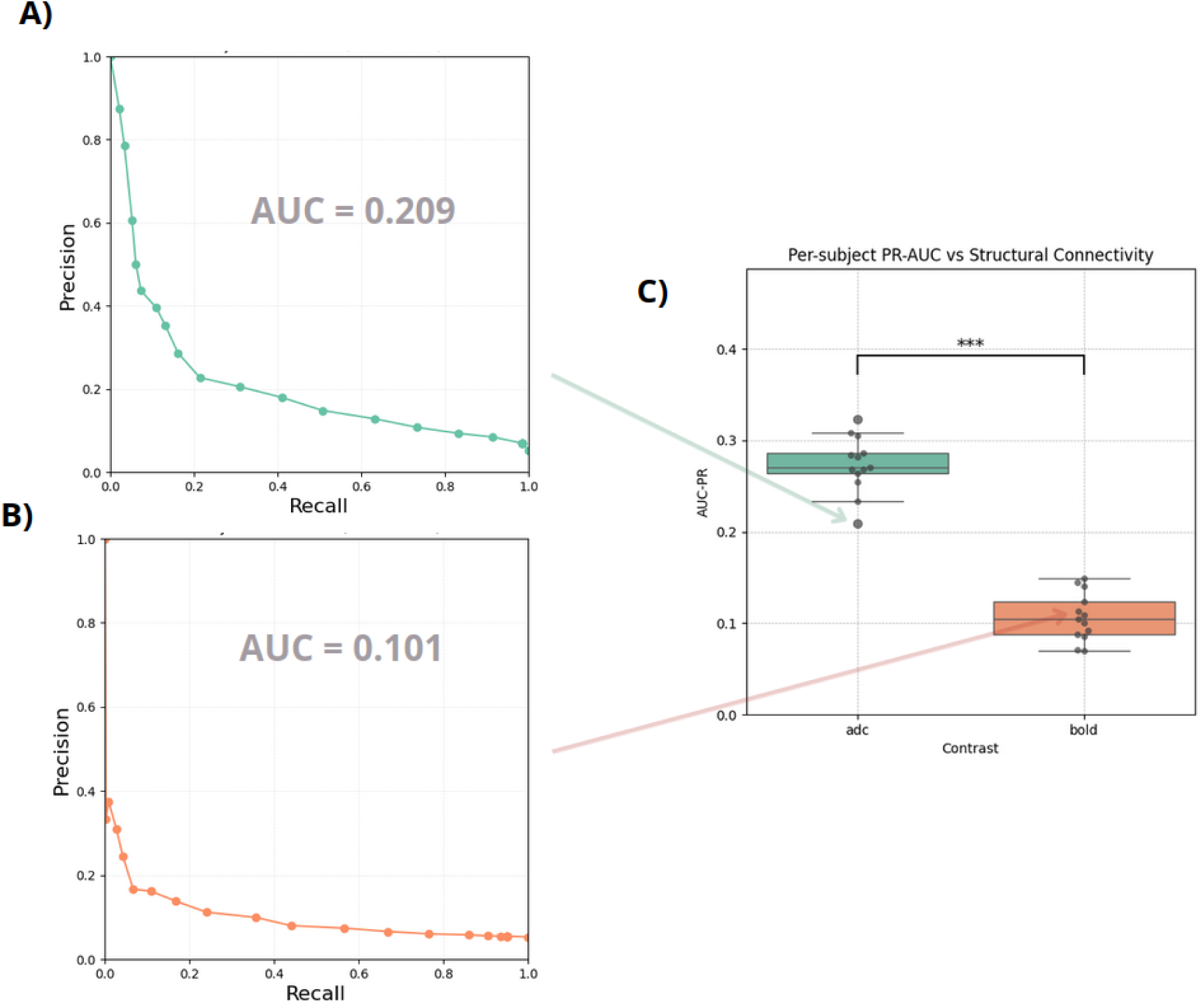
Precision-Recall (PR) Area Under the Curve (AUC) analysis. For each subject, FC matrices were binarised across a range of z-thresholds and compared against the binary SC matrix to generate PR curves for ADC and BOLD (representative subject shown in A–B). (C) Per-subject AUC-PR values for ADC (median = 0.270, interquartile range: [0.264-0.286]) and BOLD (median = 0.105, interquartile range: [0.088-0.124]). ADC showed significantly higher AUC-PR than BOLD (Wilcoxon signed-rank test, paired, two-sided; W = 0, *p* = 2.4 *×* 10*^−^*^4^, n = 13).

## 4 Discussion

### 4.1 Detecting Resting State Networks

Establishing ADC-fMRI as a reliable tool for revealing brain RSNs could further validate it as a complement to BOLD functional contrast. The fundamental distinction between these modalities lies in their underlying contrast mechanisms. Specifically, ADC is thought to measure direct neuromorphological coupling, whereas BOLD relies on an indirect neurovascular response. Despite the lower SNR of ADC-fMRI compared to BOLD [45], the partial detection of RSNs achieved here carries two important implications. First, it would confirm that the ADC timecourse SNR is nonetheless sufficient to map spontaneous ADC fluctuations effectively. Second, it lends support to the notion that those RSNs are driven by synchronous neural activity rather than being purely a product of vascular synchrony, a question still debated in the literature [79–83]. In this work, we approached RSN identification using three standard methods: ICA, seed-based correlations, and ROI-ROI FC graph analysis.

#### 4.1.1 ADC-fMRI Functional Connectivity is Predominantly Short-Ranged

In the ICA analysis, while substantial spatial overlap between subsets of BOLD and ADC independent components could be observed, the ADC components were notably more fragmented. Individual functional clusters in ADC typically appeared as isolated focal clusters, requiring manual grouping to reconstruct the matching BOLD networks. ADC’s localised topology suggests that it is limited to detecting short-range functional connections. A similar observation could be made with the seed-based analysis, which yielded much more local positive clusters than distant ones in ADC, contrasting with BOLD. Most of the few long-ranged connections detected with ADC seed-based analysis disappeared with GSR. The only ones that survived were contralateral activity within the visual and somatosensory regions.

Even when evaluating FC representations via graph analysis, ADC-fMRI yielded fewer inter-hemispheric connections than the BOLD modality. This discrepancy is fundamentally driven by the substantial anatomical distance separating these cross-hemispheric regions, further highlighting the spatial limitations of functional diffusion signals.

This dependence on distance could be attributable to the lower SNR of the ADC timecourses, which precluded the long-range connectivity seen in BOLD. Comparative maps of temporal SNR across ADC, BOLD, and dfMRI modalities revealed that ADC operates with lower temporal SNR (*≈*30) than BOLD (*≈*130) (Supplementary Figure in [45]). The data investigated in this work were acquired in the same conditions as in [45], thus these temporal SNR estimates are expected to hold. However, it must also be considered that the extended spatial reach of BOLD FC could be partially (or entirely) driven by vascular symmetry. When distant cortical regions share the same vascular supply (with the same latency), they naturally exhibit highly correlated signal fluctuations [80, 81, 83]. Compounded by systemic low-frequency oscillations travelling through the vascular tree [80, 81], these macroscopic vascular dynamics may generate false-positive long-range functional connections that do not reflect true synchronised neural firing. ADC-fMRI, on the other hand, is far less susceptible to these distal vascular contaminations [45]. Consequently, the highly localised topology observed in ADC could represent a more spatially specific mapping of local neural networks, stripped of the extended vascular blurring inherent to BOLD.

#### 4.1.2 Visual and Somatomotor Networks Show the Strongest Detection Evidence in ADC

Previous research categorises RSNs into processing and control architectures [77]. Processing networks (including the VIS and SMN) and the DMN are typically highly functionally segregated. Conversely, control networks such as the FPN and SAN are defined by their cross-network roles: the FPN serves as a dynamic flexible hub [84], while the SAN acts as a switch between the DMN and FPN [85]. Other studies differentiate RSNs along a unimodal-to-transmodal axis. Specifically, structure and function correspond closely in unimodal, (primary sensory and motor regions), but gradually untether and diverge in transmodal cortices (including the DMN) [86, 87].

Regardless of the category of each RSN, BOLD reliably resolved all six networks of interest across both independent component and seed-based analyses. Graph analysis further corroborated these detections, as every evaluated network in the BOLD modality demonstrated segregation significantly greater than a random spatial null topology. Nevertheless, the VIS, SMN, and DMN still yielded the highest segregation, aligning with expectations.

When assessing RSN detection in ADC, the VIS and SMN yielded the strongest segregation. Both networks are characterised by thick, heavily myelinated WM pathways and topographically organised short-range connectivity. These structural properties likely underpin their strong functional segregation in BOLD and, potentially, their robust detectability in ADC. Indeed, if ADC detects neural swelling, and swelling propagates most reliably along direct structural connections, then these two networks are the ideal candidates for robust ADC detection due to their strong structural tethering.

The DMN and DAN also reached group-level significance, albeit with considerably lower *z*-scores than the unimodal networks. The DMN’s detection is particularly noteworthy given longstanding debate over whether its BOLD connectivity reflects genuine neural activity or is driven by nonneural artifact [80, 81, 83, 88]. Its detection via ADC-fMRI lends support to a genuine neural basis for DMN functional organisation. Nevertheless, detection of both the DMN and DAN remained considerably less robust than that of the VIS and SMN, and this fragility was further corroborated by seed-based analyses. While seed-based methods successfully recovered bilateral FC within the SMN and VIS that proved resilient to GSR, the DMN and DAN displayed substantially weaker FC, with initial positive clusters only partially overlapping with expected anatomical boundaries before being entirely abolished following GSR. Interestingly, for the DMN, we observe positive clusters outside of the expected areas, with positive clusters emerging in WM connecting the precuneus to the angular gyri and to the frontal lobe, suggesting functional coupling along these structural pathways [88]. Also, GSR unmasked sparse negative clusters in VIS, SMN and DMN networks that spatially converged with anticipated anticorrelated ROIs, mirroring patterns observed in the BOLD data.

In contrast, the FPN and SAN did not survive significance testing. These two networks are distinguished by their role as flexible integrative hubs that dynamically reconfigure across cognitive states [84, 89], resulting in inherently lower static temporal cohesion when averaged across a resting-state scan. Consequently, while BOLD captured connectivity within these transmodal systems, ADC appeared selectively sensitive to networks with more robust static functional cohesion.

Taken together, these results suggest a gradient of ADC detectability shaped by two partially overlapping properties: static functional cohesion and structural tethering. The VIS and SMN score highly on both dimensions, yielding the most robust detection. The DMN and DAN are sufficiently segregated for detection in graph analysis, but their weaker structural grounding renders this detection fragile across analytical approaches. The FPN and SAN, as the most dynamically reconfiguring networks, lack the static cohesion necessary for ADC detection altogether. Whether these three tiers reflect a true hierarchy of structural tethering, temporal stability, or a combination of both remains an open question that future work with larger cohorts and complementary structural imaging could address.

### 4.2 Including WM in FC

#### 4.2.1 Enhanced Detection of Grey-White Matter Coupling via the ADC

Having established which RSNs ADC-fMRI can resolve in GM, we next investigate whether its tissue-agnostic contrast mechanism gives it an advantage when connectivity spans the GM-WM boundary. Our results reveal a distinct divergence in modality sensitivity: while BOLD-fMRI identified a higher number of significant GM-GM edges, ADC-fMRI proved more sensitive to functional links involving the WM. This suggests that while BOLD remains highly effective for mapping cortical-cortical interactions, ADC-fMRI may offer a complementary advantage for resolving FC within the less vascularized environment of WM tracts [41–44].

The reduced sensitivity of BOLD to connectivity to WM in our analysis directly reflects the well-documented haemodynamic bottlenecks of these pathways. The attenuation we observed aligns with the tissue’s inherently sparse vascularity [28]. More critically, the reliable detection of synchronous BOLD activity across regions is disrupted by the depth-dependent variability of the WM HRF [29]. Even when temporal correlations are found, interpreting them as functional links remains problematic due to conflicting evidence regarding their biological origin, specifically, whether these vascular fluctuations reflect genuine neural signalling [20] or merely isolated glial metabolic maintenance [31]. This ongoing physiological ambiguity makes the inclusion of WM in BOLD-derived FC analyses rather controversial.

In contrast, ADC-fMRI provides a more uniform response across different tissue types by bypassing the complex and contentious neurovascular coupling that governs the BOLD signal. By relying on a mechanism that appears less dependent on local vascular architecture, ADC-fMRI avoids the haemodynamic discrepancies that typically challenge correlations between high-perfusion GM and low-perfusion white matter. Consequently, ADC-fMRI proves to be an effective tool for bridging these tissue types, providing a more direct proxy for neural synchrony within the WM [39, 41–44].

#### 4.2.2 ADC Exhibits Stronger Structure-Function Alignment than BOLD

Following the observation that ADC was more sensitive to GM-WM connectivity than BOLD, we conducted a further investigation by plotting FC graphs restricted to significant GM-WM links. We noted that certain WM nodes exhibited extensive connections to GM regions in BOLD, while others showed none at all, as previously observed in [18, 20]. Furthermore, the BOLD data revealed significant connectivity between association fibres in one hemisphere and GM regions in the contralateral hemisphere, despite the absence of direct structural links. This phenomenon was far less prevalent in the ADC results, where WM nodes that correlated with GM nodes in both hemispheres were predominantly identified as commissural fibres. Further analysis using PR-AUC demonstrated that a higher alignment could be measured between ADC and structural connectivity than BOLD and structure.

At first glance, our standard BOLD results may appear at odds with previous literature that has successfully demonstrated structure-function coupling between WM BOLD signals and GM regions [18–20, 25]. Such coupling has been documented in the RS, during task performance, and across pathological conditions [21, 23]. However, resolving this apparent inconsistency of our results compared to previous work highlights the core methodological advantage of ADC-fMRI. These previous studies explicitly acknowledge that the WM BOLD signal is highly confounded by tissue composition, microstructure, and the physiological ambiguities discussed above. To extract meaningful structure-function alignment, recent work has been forced to abandon traditional Pearson’s correlations in favour of highly sophisticated analytical frameworks. These include methods enabling the assessment of WM engagement in GM connectivity [22, 24], mediation analyses [24], and dynamic FC approaches that correlate GM pairs with the windowed variance of the connecting tract [26].

What our study demonstrates is that these complex analytical interventions are unnecessary when utilising ADC-fMRI. By inherently bypassing the haemodynamic variability of WM signals, this modality yields GM-WM FC that demonstrates superior alignment with underlying SC, using standard Pearson correlations alone. Future work could certainly apply these advanced BOLD frameworks to our dataset to see if they close the performance gap, but the present results underscore ADC as a potentially more direct tool for mapping whole-brain connectivity.

## 5 Limitations

A primary limitation of this study stems from the fundamental acquisition scheme used to compute the ADC timeseries. As previously outlined, each single ADC time point is derived from a pair of diffusion-weighted volumes acquired sequentially with different b-values. Consequently, if the brain undergoes a functional state transition between these two successive acquisitions, the resulting ADC calculation effectively conflates these distinct states into a single, averaged data point. This temporal blurring significantly restricts the ability to capture highly dynamic RSNs.

Furthermore, as demonstrated during the ICA denoising procedure, the dMRI data were susceptible to slice cross-talk artifacts resulting from the multiband acceleration factor of 3. Specific independent components exhibited distinct spatial patterns characterised by high-intensity signal fluctuations restricted entirely to the three simultaneously acquired slices. Although efforts were made to identify and regress out these artefactual components, it is possible that residual slice leakage persisted in the data. This residual artifact carries the risk of artificially inflating FC estimates between spatially distant regions that happen to share an acquisition slice group.

The sample size of this study was limited to 13 subjects, which constrains the statistical power of the analyses and the generalisability of the findings. Nevertheless, despite this modest cohort, ADC-fMRI successfully detected 4 out of 6 RSNs using graph analysis, suggesting that the reported effects are robust enough to emerge even under such conditions. Replication in larger cohorts will be necessary to confirm these findings, particularly for the DMN and DAN, whose detection was less consistent across analytical approaches.

The seed-based analysis for detecting RSNs relied on the a priori selection of a single, representative seed region per network. Exhaustively testing every possible ROI as a seed for each network would have been computationally prohibitive and practically difficult. Consequently, while this targeted approach was necessary, FC remains inherently sensitive to seed placement, and alternative selections could yield variations in the observed network topographies. Second, the graph analysis defined significant segregation effects using an assigned threshold of *Z >* 2.3. As with any predetermined statistical threshold, applying a more stringent criterion could alter the findings, potentially reducing the detectability of the DMN for example.

In the context of the GM-WM connectivity analysis, a notable limitation is the reliance on a population-based probabilistic structural connectivity atlas (derived from 66 subjects) rather than subject-specific tractography. Because these normative WM ROIs do not account for individual anatomical variations in tract trajectories, there is an increased risk of spatial misalignment between the standard template and the subjects’ true native anatomy. Such discrepancies may have introduced variance into the FC-SC coupling estimates.

## 6 Conclusions

This study investigated FC by applying analytical methods conventionally used for BOLD-fMRI RSNs to ADC-fMRI, with the dual aim of characterising ADC’s sensitivity to cortical RSNs and its ability to capture GM-WM functional coupling. This comparative approach highlights key modality differences, particularly concerning their dependence on vascular dynamics and their inherent sensitivity to neural activity across tissue types.

Group-level ICA of ADC data yielded spatially constrained components requiring manual aggregation to reconstruct RSNs comparable to those observed in BOLD, and seed-based analyses predominantly identified clusters in close proximity to the seed region. Whether this spatial confinement reflects a sensitivity limitation of ADC or whether BOLD artificially merges distinct neural regions sharing common vascular supplies remains an open question. Despite this proximal bias, seed-based analyses successfully recovered robust contralateral connectivity within the SMN and VIS networks. ROI-ROI graph analysis revealed a gradient of ADC detectability across the six evaluated networks. The VIS and SMN, being both highly functionally segregated and strongly tethered to underlying structural pathways, yielded the most robust detection.

The DMN and DAN reached group-level significance albeit with lower *z*-scores, and their detection proved fragile in seed-based analyses, suggesting that functional segregation alone may be insufficient for cross-analytical stability without strong structural grounding. The FPN and SAN, whose functional identity is defined by dynamic cross-network reconfiguration rather than internal cohesion, did not reach significance. This gradient of detectability suggests that ADC-fMRI is selectively sensitive to networks with robust static functional cohesion and structural tethering, offering a window into the neural rather than vascular substrate of RSN organisation.

Finally, targeted investigations into GM-WM interactions demonstrated that ADC-fMRI is arguably superior to BOLD for such analyses, with GM-WM FC showing stronger alignment with underlying structural connectivity across subjects. This advantage likely reflects ADC’s tissue-agnostic contrast mechanism, which captures functional signals in white matter without the vascular confounds that limit BOLD in these regions.

In summary, these findings advocate for ADC-fMRI as a complementary modality to BOLD. Its decoupling of functional signals from vascular haemodynamics provides critical insights into the structural and neural foundations of brain connectivity that BOLD cannot access directly.

## 8 Data and Code Availability

Raw data are available at https://doi.org/10.5281/zenodo.17590877. The code will be released upon the publication of the article.

## 9 Author Contribution

Conceptualisation: I.J, J.N.D; Methodology: I.J, J.N.D; Investigation: A.S., J.B.P., J.N.D.; Data curation: A.S., J.B.P., J.N.D.; Validation: J.N.D; Formal analysis: J.N.D; Writing-original draft: J.N.D; Writing - Review & Editing: J.N.D, I.J, A.S, I.d.R, J.B.P, S.A., T.P.; Visualisation: J.N.D; Supervision: I.J.;

## 10 Acknowledgements

This work was supported by the Swiss Secretariat for Research and Innovation (SERI) under an ERC Starting Grant award ‘FIREPATH’ MB22.00032, and by Swiss National Science Foundation (SNSF) grants no. 194260 and 10000465. During the preparation of this work, Claude Sonnet 4.6 was used to assist with the grammatical refinement of the paper. We thank Filip Szczepankiewicz for his contribution to this work through the development of the isotropic waveform for ADC-fMRI.

