## Supplementary Materials for "A Comprehensive Analysis Comparing Isotropic ADC to BOLD-fMRI: Sensitivity to Resting State Networks and Grey to White Matter Functional Connectivity"

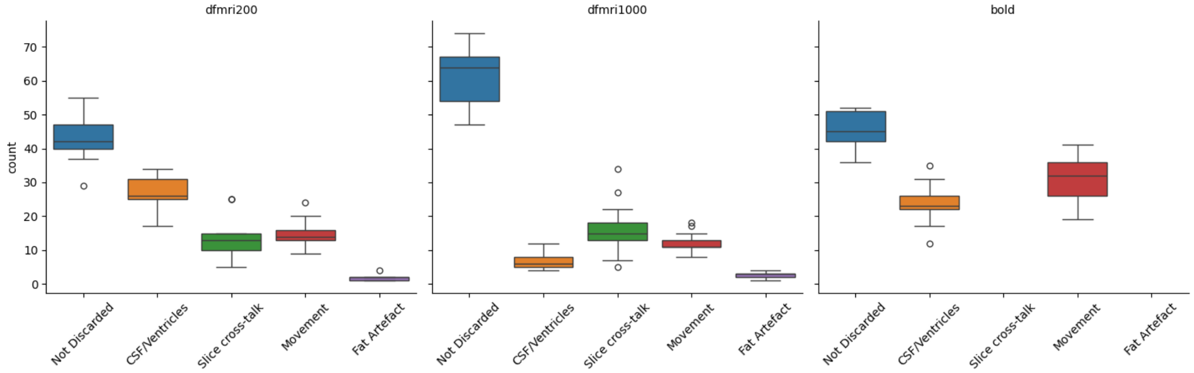


Figure S1: ICA cleaning procedure: for each contrast, the number of independent components (out of 100) that were retained (not discarded), as well as those discarded due to CSF/ventricular signal, slice cross-talk, motion, and fat artefacts, is shown. Discarded components were regressed out of the timeseries, and the observed variability reflects inter-subject differences.

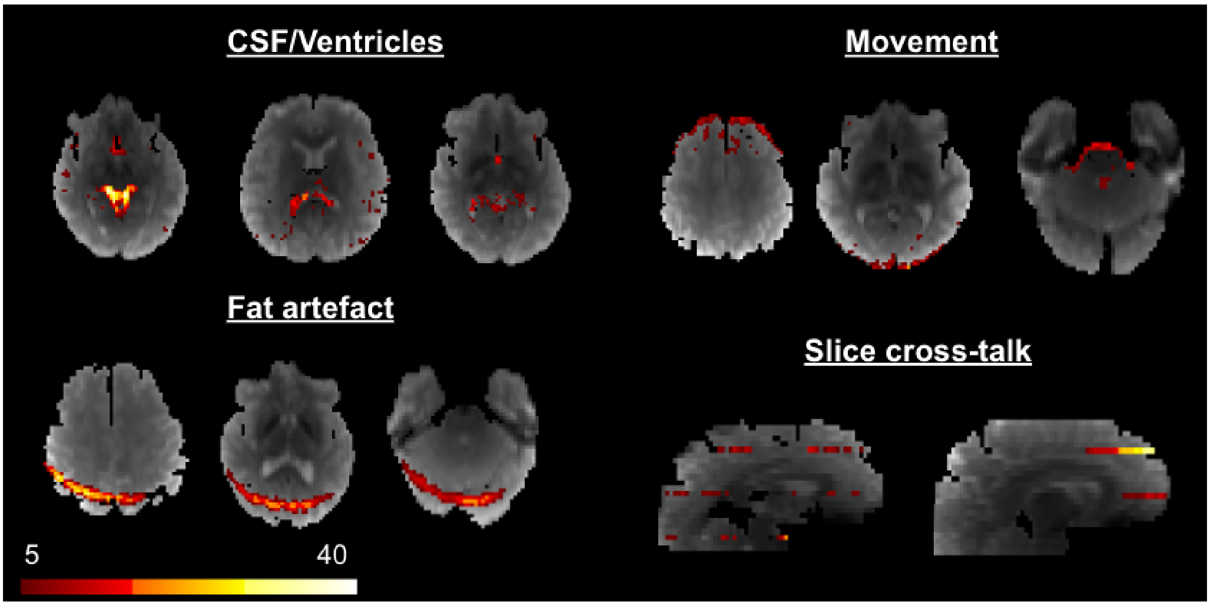


Figure S2: Examples of categories of regressed independent components in an example subject.

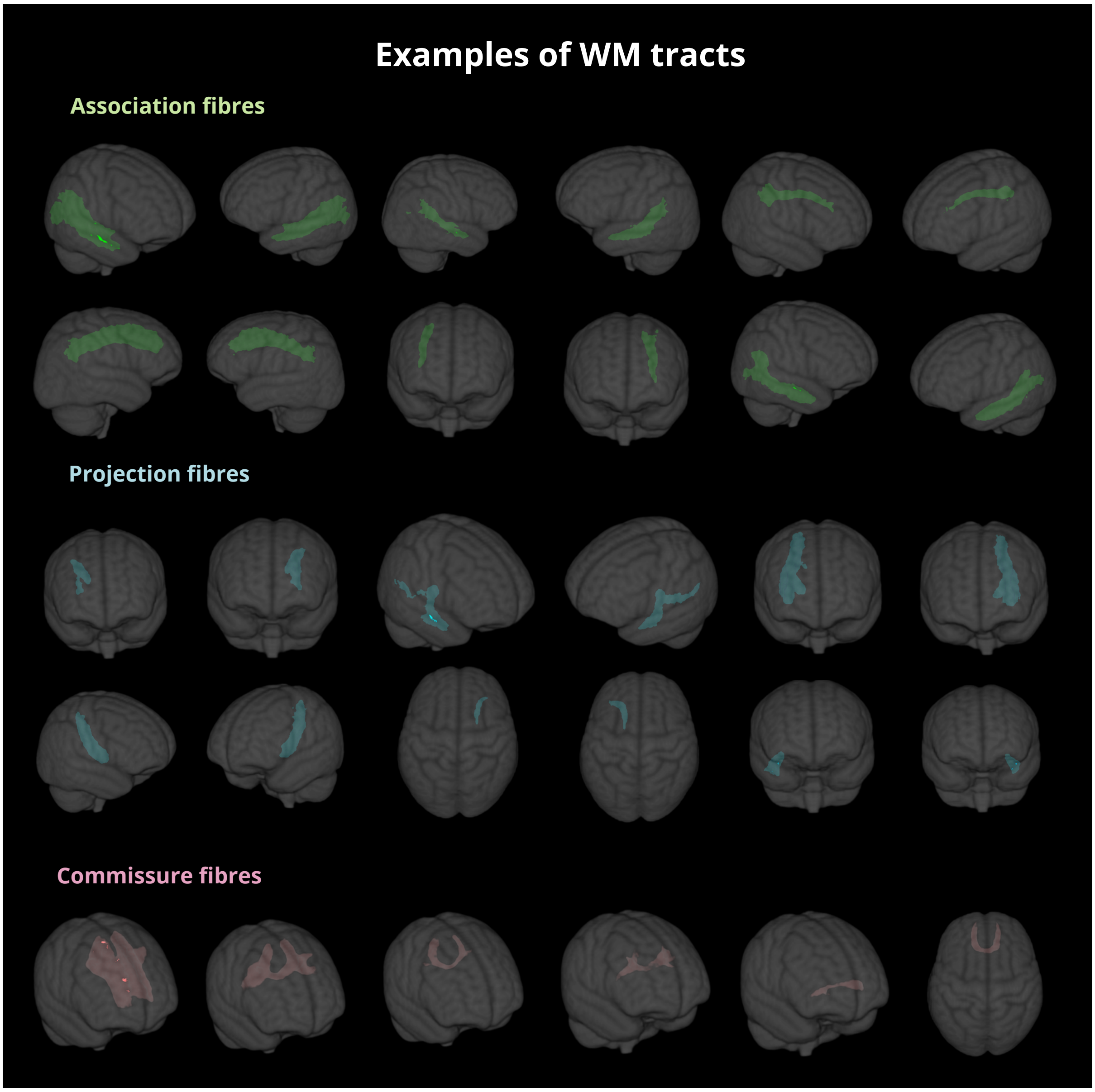


Figure S3: Overview of examples of WM fibres. Association, Projection and Commissural fibres were obtained by combining highly overlapping WM tracts obtained from the *probconnatlas* tool [1]. In total, 90 WM fibres were retained.

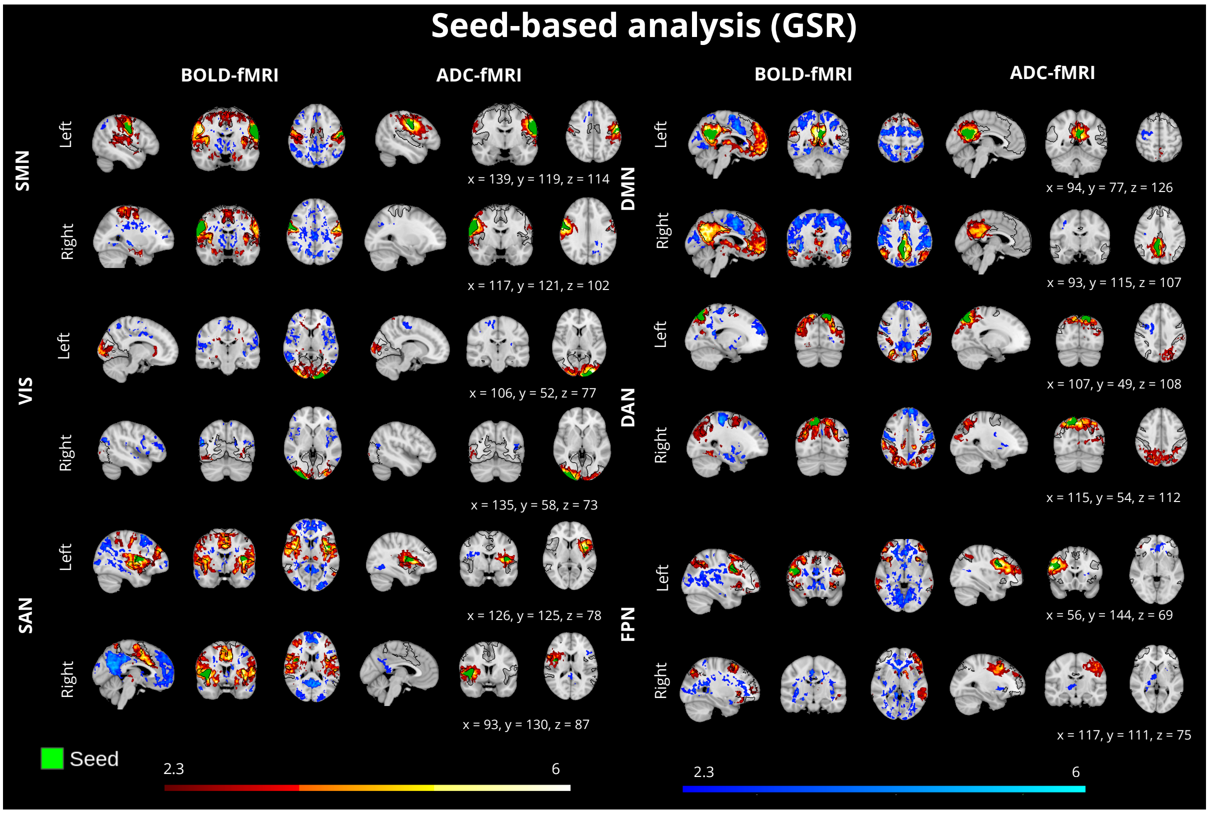


Figure S4: Group-level seed-based analysis performed with GSR on seeds including: the precuneus/posterior cingulate (DMN), lateral prefrontal cortex (FPN), precentral gyrus (SMN), lateral occipital cortex (VIS), superior parietal cortex (DAN), and anterior insula/operculum (SAN). Statistical maps were obtained using FSL FEAT with mixed-effects modelling (FLAME 1+2) and thresholded at an absolute $z > 2.3$ (cluster-corrected) to identify significant voxels.
